# Genetic interactions between the chromosome axis-associated protein Hop1 and homologous recombination determinants in *Schizosaccharomyces pombe*

**DOI:** 10.1101/190439

**Authors:** Simon David Brown, Olga Dorota Jarosinska, Alexander Lorenz

## Abstract

Hop1 is a component of the meiosis-specific chromosome axis and belongs to the evolutionarily conserved family of HORMA domain proteins. Hop1 and its orthologs in higher eukaryotes are a major factor in promoting double-strand DNA break formation and inter-homolog recombination. In budding yeast and mammals they are also involved in a meiotic checkpoint kinase cascade monitoring the completion of double-strand DNA break repair. We used the fission yeast, *Schizosaccharomyces pombe*, which lacks a canonical synaptonemal complex to test whether Hop1 has a role beyond supporting the generation of double-strand DNA breaks and facilitating inter-homolog recombination events. We determined how mutants of homologous recombination factors genetically interact with *hop1*, studied the role(s) of the HORMA domain of Hop1, and characterized a bio-informatically predicted interactor of Hop1, Aho1 (SPAC688.03c). Our observations indicate that in fission yeast, Hop1 does require its HORMA domain to support wild-type levels of meiotic recombination and localization to meiotic chromatin. Furthermore, we show that *hop1Δ* only weakly interacts genetically with mutants of homologous recombination factors, and in fission yeast likely has no major role beyond break formation and promoting inter-homolog events. We speculate that after the evolutionary loss of the synaptonemal complex, Hop1 likely has become less important for modulating recombination outcome during meiosis in fission yeast, and that this led to a concurrent rewiring of genetic pathways controlling meiotic recombination.

## Introduction

Chromosomes of diploid organisms are present in two sets, each derived from one of the parents. To reproduce sexually, this genetic material is halved exactly to produce haploid cells that contain only one chromosome set, the gametes. During fertilization, two gametes from different sexes fuse to form a zygote, re-establishing a double chromosome set. A specialized kind of cell division, called meiosis, is employed to halve the genetic material. In meiosis, one round of chromosome duplication is followed by two rounds of chromosome segregation. Remarkably, correct segregation of chromosomes in meiosis requires formation of double-strand DNA breaks (DSBs) by the conserved transesterase Spo11 (Lam and Keeney 2015), followed by repair through homologous recombination. Homologous recombination, in combination with sister chromatid cohesion, establishes physical connections (chiasmata) between maternal and paternal chromosomes that guide their proper segregation, and promotes reciprocal exchange (crossovers) of maternal and paternal genetic information (Hunter 2015). Careful regulation of this complex process is imperative to ensure the required number of physical links between the correct partner chromosomes, and that unrepaired DNA breaks are mended (Lam and Keeney 2015; Hunter 2015). The repair of the Spo11-induced DSBs is started by DNA resection. During the initial step of DNA resection Spo11, which is covalently linked to the DNA ends, is endonucleolytically removed by the Mre11-Rad50-Nbs1 (Mre11-Rad50-Xrs2 in *Saccharomyces cerevisiae*) complex in cooperation with Ctp1 (Sae2 in *S. cerevisiae*) (Neale et al. 2005; Hartsuiker et al. 2009). Long-range resection then exposes substantial 3’ tails which are used to invade homologous template DNA on the homologous chromosome or the sister chromatid (strand exchange) to mend the DSB by recombination (reviewed in Mimitou and Symington 2011; Hunter 2015; Lorenz 2017). Key factors for the strand exchange process in meiosis are the RecA-type recombinases Rad51 and Dmc1 (Shinohara et al. 1997; Grishchuk and Kohli 2003) supported by their mediators Rad55-Rad57, Swi5-Sfr1, and Rlp1-Rdl1-Sws1 (SHU complex in *S. cerevisiae*) (Tsutsui et al. 2001; Grishchuk and Kohli 2003; Akamatsu et al. 2003; Martín et al. 2006; Sasanuma et al. 2013; Hong et al. 2013; Lorenz et al. 2014). These Rad51/Dmc1-mediators are important for generating and stabilizing Rad51/Dmc1 coated nucleoprotein filaments which then invade homologous template DNA (Sung 1997; Haruta et al. 2006). The DNA on the homologous chromosome (or sister chromatid) being invaded is opened as a displacement loop (D-loop) which can then be extended by DNA synthesis using the invading sequence as primer (reviewed in Brown and Bishop 2014). The way in which D-loops or recombination intermediates derived from it are processed, determines whether repair will occur as a crossover (CO) or a non-CO, and Rad51/Dmc1-mediators play an important role in this (reviewed in Brown and Bishop 2014; Hunter 2015; Lorenz 2017).

In fission yeast processing of most meiotic recombination intermediates into COs and some non-COs depends on the structure-selective endonuclease Mus81-Eme1, the lack of which results in meiotic catastrophe and strongly reduced spore viability (Osman et al. 2003; Smith et al. 2003). Additionally, a sizeable fraction of non-COs is produced by the action of the FANCM-ortholog Fml1, a DEAD/DEAH-box DNA helicase, and it has been shown that Mus81-Eme1 and Fml1 work in parallel to process meiotic recombination intermediates (Lorenz et al. 2012). In a wild-type situation Fml1 action seems to be curtailed by Rad51/Dmc1-mediators, because weakening the strand exchange reaction by removing any Rad51/Dmc1-mediator in a *mus81Δ* background ameliorates the strong spore viability defect of *mus81Δ* without restoring CO formation; this rescue depends on the presence of Fml1 (Lorenz et al. 2012, 2014).

To facilitate the generation of DSBs and their correct repair during meiosis, chromosomes undergo a progression of changes starting with the establishment of meiosis-specific chromosome axes and, in most organisms, culminating in the formation of the synaptonemal complex „ an elaborate proteinaceous structure (reviewed in Cahoon and Hawley 2016). Components of the synaptonemal complex seem to be critical for controlling several aspects of meiotic chromosome metabolism, from pairing of homologous chromosomes, to initiation of recombination and maturation of the physical interactions between the homologous chromosomes (reviewed in Cahoon and Hawley 2016; Gray and Cohen 2016). However, some organisms have overcome the necessity for a fully-fledged synaptonemal complex and perform meiosis with a simplified version of it (reviewed in Loidl 2016). The fission yeast *Schizosaccharomyces pombe* is the best-studied organism executing meiosis without a canonical synaptonemal complex (reviewed in Loidl 2006).

Despite lacking a synaptonemal complex, fission yeast meiotic chromosomes assemble axes, called linear elements (Olson et al. 1978; Bähler et al. 1993; Molnar 2003), consisting of proteins which are evolutionarily related to axial/lateral elements of the synaptonemal complex (Lorenz et al. 2004), and building upon the existing mitotic chromosome axis organization (reviewed in Ding et al. 2016; Iwasaki and Noma 2016). One of these components is Hop1 which, in contrast to other meiotic chromosome axis factors, is conserved on the protein sequence level with homologs from yeast to man (Lorenz et al. 2004; Rosenberg and Corbett 2015). In both *S. cerevisiae* and *Sz. pombe*, Hop1 localizes to the meiosis-specific chromosome axis (Hollingsworth et al. 1990; Smith and Roeder 1997; Lorenz et al. 2004), and deletion of *HOP1/hop1^+^* causes a decrease in meiotic recombination and homologous chromosome pairing (Hollingsworth and Byers 1989; Loidl et al. 1994; Latypov et al. 2010). Hop1 is characterized by a conserved N-terminal HORMA (<u>Ho</u>p1-<u>R</u>ev7-<u>Ma</u>d2) domain (Aravind and Koonin 1998) (Fig. 1a) and is the founding member of the meiotic HORMA domain (HORMAD) family proteins. In contrast to yeasts which contain only one meiotic HORMAD (Hop1), higher eukaryotes contain multiple paralogs of meiotic HORMADs: These include HIM-3, HTP-1, HTP-2, and HTP-3 in *C. elegans*; ASY1 and ASY2 in *Arabidopsis*; and HORMAD1 and HORMAD2 in mammals (Zetka et al. 1999; Caryl et al. 2000; Couteau and Zetka 2005; Martinez-Perez and Villeneuve 2005; Goodyer et al. 2008; Fukuda et al. 2010). In fungi, plants, and mammals the HORMA domain is followed by several [S/T]Q residues representing potential Tel1/ATM-Rad3/Mec1/ATR phosphorylation sites, which enable meiotic HORMADs to act as structural adaptors for a meiotic checkpoint kinase cascade (Sanchez-Moran et al. 2007; Carballo et al. 2008; Daniel et al. 2011; Kogo et al. 2012; Osman et al. 2018) (Fig. 1a). Fungal Hop1 proteins also contain a centrally located CxxC Zn-finger motif (Hollingsworth et al. 1990) (Fig. 1a) which has been demonstrated to be important for *in vitro* DNA-binding of *S. cerevisiae* Hop1 (Tripathi et al. 2007). Despite its *in vitro* DNA-binding capabilities, Hop1 requires the presence of Rec10 (Red1 in *S. cerevisiae*) to localize to chromosome axes *in vivo* (Smith and Roeder 1997; Lorenz et al. 2004); indeed, Hop1 and Rec10/Red1 have been shown to physically interact with each other (de los Santos and Hollingsworth 1999; Spirek et al. 2010). Rec10 in turn is recruited to chromosome axes by the meiosis-specific cohesin subunit Rec11 upon phosphorylation by casein kinase 1 (Phadnis et al. 2015; Sakuno and Watanabe 2015).

**Figure 1.**
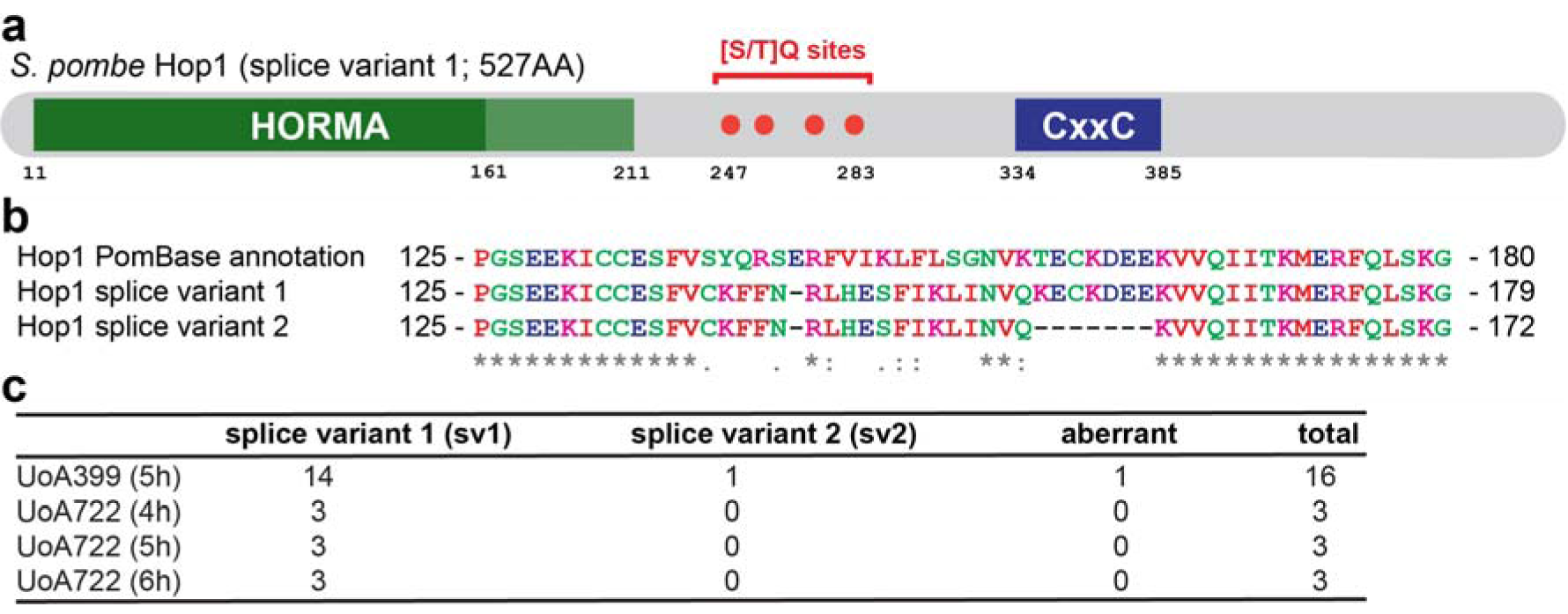
*Schizosaccharomyces pombe* Hop1 domain structure and alignment between annotated and experimentally determined versions. (**a**) Schematic of the domain structure of the experimentally derived sv1 (splice variant 1) version of Hop1 with the HORMA domain at the N terminus (AAs 11-211), four potential [S/T]Q phosphorylation sites (red dots) distributed between AAs 247 to 283, and residues 334-385 comprising the CxxC Zn-finger domain (dark blue). The HORMA domain can be subdivided into a core domain (dark green) represented by the most C-terminal ~150 AAs and a “safety-belt” region (light green; residues 161-211) (Rosenberg and Corbett 2015; West et al. 2017). Positions of key AA residues are indicated below the schematic. (**b**) ClustalΩ alignment (Sievers et al. 2014) of part of the HORMA domain (starting at residue 125) of the current PomBase-annotation of Hop1, and the two observed splice variants Hop1-sv1 and Hop1-sv2. Colouring of AAs follows the standard ClustalΩ scheme according to physicochemical properties: red for small and small & hydrophobic AAs (AVFPMILW); blue for acidic AAs (DE); magenta for basic AAs (RK); and green for AAs containing hydroxyl-, sulfhydryl-, or amine-groups. Consensus symbols below the alignment indicate full conservation (*) and conservation of groups with strongly similar (:) or weakly similar (.) properties, for details see http://www.ebi.ac.uk/Tools/msa/clustalo/help/faq.html. (**c**) Relative abundance of different hop1 splice variants found in cDNA of the indicated diploid strains at given time-points of a meiotic time-course.

Here, we investigate genetic interactions of *hop1* with meiotic recombination determinants in fission yeast to test whether *hop1* has (a) role(s) beyond its established function in promoting DSB formation (Latypov et al. 2010). Genetic interactions with factors important for recombination intermediate processing (*mus81* and *fml1*) suggested a potential, likely indirect, role for Hop1 in the strand exchange process. However, subsequent epistasis analysis with the strand exchange factor *dmc1* and Rad51/Dmc1-mediators (*rad55, sfr1*, and *rlp1*) did not support the hypothesized separate function of Hop1 in strand exchange. Furthermore, we also show that the HORMA domain is required for wild-type levels of meiotic recombination, and for localization to meiotic chromatin in *Sz. pombe*. Implications of the evolutionary differences regarding meiotic HORMAD proteins and their interactions are discussed.

## Material and methods

### *Sz. pombe* and *E. coli* culture conditions

*E. coli* strains were grown on LB and SOC media „ where appropriate containing 100 µg/ml Ampicillin (Sambrook and Russell 2000). Competent cells of *E. coli* strains NEB10^^®^^-beta (New England BioLabs Inc., Ipswich, MA, USA), and XL1-blue (Agilent Technologies, Santa Clara, CA, USA) were transformed following the protocols provided by the manufacturers.

*Sz. pombe* strains used for this study are listed in Supplementary Table S1. Yeast cells were cultured on yeast extract (YE), yeast nitrogen base glutamate (YNG) and Edinburgh minimal media with glutamate as nitrogen source (EMMG) agar plates containing the required supplements (concentration 250 mg/l on YE, 75 mg/l on YNG and EMMG). Crosses were performed on malt extract (ME) agar containing the required supplements at a final concentration of 50 mg/l, unless strains to be crossed contained plasmids, in which case they were sporulated on sporulation agar with supplements (SPAS) plates with the necessary supplements at 50 mg/l to maintain selection for the plasmids (Smith 2009; Sabatinos and Forsburg 2010). Transformation of yeast strains was done using a standard lithium-acetate procedure (Keeney and Boeke 1994) with the modifications described in Brown and Lorenz (2016).

To obtain meiotic *Sz. pombe* cells for chromatin spreading and cDNA preparation meiotic time-courses were executed according to a published protocol (Loidl and Lorenz 2009).

### Yeast strain and plasmid construction

For specifics of strain and plasmid construction, please refer to Supplementary Materials.

Construction of the *kanMX6*-marked *hop1Δ −1* allele (Latypov et al. 2010) and the kanMX6-marked *mug20^+^*-GFP (Estreicher et al. 2012) have been described previously. The *natMX6*-marked *hop1Δ −25* was derived from *hop1Δ −1* using an established marker swap protocol (Sato et al. 2005; Lorenz 2015).

To produce *hop1*-alleles expressing a particular splice variant or the *hop1-ΔHORMAD* mutant, we first partially deleted *hop1* retaining the coding sequence directly upstream of the first and directly downstream of the last intron using a *ura4* marker (Grimm et al. 1988). The resulting strain carrying the *hop1-3* allele (UoA725) was subsequently transformed with the cDNA of one of the two splice variants or a *hop1-ΔHORMAD* construct; candidate strains were selected by resistance to FOA (5-fluoroorotic acid) and then subjected to further testing.

Epitope-tagging of Hop1 with 13myc was achieved by cloning a transformation cassette into pFA6a-*13myc-kanMX6* (Bähler et al. 1998) with sequences from the 3’ end and downstream flanking region of the *hop1* coding sequence. This transformation cassette was amplified by PCR and transformed into strains UoA746, UoA868, and UoA938.

A deletion cassette for aho1 was constructed by cloning flanking sequences of *SPAC688.03c* into pAG25 up- and downstream of the *natMX4* marker (Goldstein and McCusker 1999). The resulting plasmid (pALo116) was linearized by a restriction digest and transformed into the standard lab strain FO652.

To over-express *aho1^+^*, the coding sequence of SPAC688.03c was amplified by PCR and cloned into pREP41-*eGFPC* (Craven et al. 1998).

To construct plasmids for over-expression of *hop1^+^* in *Sz. pombe* we cloned PCR products representing genomic or cDNA versions of the *hop1* coding sequence into pJR-41XL (Moreno et al. 2000). During DNA sequencing we found discrepancies with the *hop1* coding sequence prediction on PomBase (https://www.pombase.org/, last accessed 03/06/2017) (see Results section for details). To corroborate this initial finding we also cloned meiotic cDNA clones from two independently constructed strains (UoA399 and UoA722) into pUC8.

All plasmid constructs (for sequences see supporting online material, Lorenz 2018), as well as epitope-tagged, splice variant, and *hop1-ΔHORMAD* strains were verified by DNA sequencing (Source BioScience plc, Nottingham, UK).

All DNA modifying enzymes (high-fidelity DNA polymerase Q5, restriction endonucleases, T4 DNA ligase) and NEBuilder HiFi DNA Assembly Master Mix were obtained from New England BioLabs. Oligonucleotides were supplied by Sigma-Aldrich Co. (St. Louis, MO, USA).

### Genetic and molecular biology techniques

Spot assays to monitor growth and genotoxin sensitivities were essentially performed as described previously (Doe et al. 2000; Lorenz et al. 2009). Spot assays of deletion strains were done on YE plates, and of strains harboring plasmids on EMMG plates containing the required supplements. To repress gene expression from pREP41-type plasmids in strains grown on EMMG, thiamine was added to a final concentration of 4 μM (Sabatinos and Forsburg 2010).

Genetic recombination assays and spore viability testing by random spore analysis have been described in detail (Osman et al. 2003; Lorenz et al. 2010, 2012, 2014; Sabatinos and Forsburg 2010).

Meiotic cDNA was obtained from cells undergoing synchronized azygotic wild-type meiosis. Aliquots of cell culture were drawn at several time-points during a meiotic time-course by first extracting RNA using the NucleoSpin TriPrep kit (Macherey-Nagel GmbH & Co. KG, Düren, Germany) and then reverse-transcribing the RNA into cDNA using the ProtoScript II First Strand cDNA Synthesis kit (New England BioLabs) following the instructions of the respective manufacturer.

### Chromatin spreads, immunocytochemistry, and microscopy

Nuclear spreads of meiotic *Sz. pombe* cells were prepared as described previously (Loidl and Lorenz 2009), with the only modification that Lallzyme MMX (Lallemand Inc., Montréal, Canada) at a final concentration of 100 mg/ml was used as the sole enzyme in the spheroplasting solution (Flor-Parra et al. 2014).

Immunostaining was performed as previously described (Loidl and Lorenz 2009) using polyclonal rabbit α-myc (ab9106; Abcam PLC, Cambridge, UK) at a 1:500 dilution and monoclonal rat α-GFP [3H9] (ChromoTek GmbH, Planegg-Martinsried, Germany) at a 1:100 dilution as primary antibodies. Antibody-bound protein was visualized using donkey α-rabbit IgG AlexaFluor-555 (ab150062; Abcam) and donkey α-rat IgG AlexaFluor-488 (ab150153; Abcam), both at a 1:500 dilution, as secondary antibodies conjugated to fluorophores. DNA was stained by Hoechst 33342 (Molecular Probes, Eugene, OR, USA) at a final concentration of 1 μg/ml.

Analysis was done using a Zeiss Axio Imager.M2 (Carl Zeiss AG, Oberkochen, Germany) epifluorescence microscope equipped with the appropriate filter sets to detect red, green, and blue fluorescence. Black-and-white images were taken with a Zeiss AxioCam MRm CCD camera controlled by AxioVision 40 software v4.8.2.0. Images were pseudo-coloured and overlayed using Adobe Photoshop CC (Adobe Systems Inc., San José, CA, USA).

### Data presentation and statistics

Data presented as box-and-whisker plots were created in RStudio 1.0.136 (R version 3.3.1) (http://www.r-project.org/) using the boxplot() function with its standard settings. The lower and upper ‘hinges’ of the box represent the first and third quartile, respectively, and the black bar within the box indicates the median (=second quartile). The ‘whiskers’ represent the minimum and maximum of the range, unless they differ more than 1.5-times the interquartile distance from the median. In the latter case, the borders of the 1.5-times interquartile distance around the median are indicated by the ‘whiskers’ and values outside this range (‘outliers’) are shown as open circles. Raw data and R scripts used for the boxplot() function are available online as supporting material (Lorenz 2018).

Initially, all data involving multiple comparisons underwent an overall statistical assessment using a Kruskal-Wallis one-way analysis of variance to verify that there indeed are statistically significant differences (α < 0.05) between data sets within one experiment. All direct one-on-one statistical comparisons between control and experimental data were done using a two-tailed Mann-Whitney U test. Both Kruskal-Wallis and Mann-Whitney U tests are non-parametric and therefore do not assume normal distribution of data points within a data set. Kruskal-Wallis analyses were performed in IBM SPSS Statistics 24.0.0.0 (International Business Machines Corporation, Armonk, NY, USA), and Mann-Whitney U tests in G*Power 3.1.9.2 (Faul et al. 2007, 2009). P values listed in Supplementary Tables S2 and S3 were determined with a statistical power (1 – β) set to 0.8.

## Results

### Correction of the *Sz. pombe* hop1 coding sequence

When constructing plasmids to over-express *hop1^+^* in fission yeast cells, sequencing of our cDNA clones revealed major discrepancies compared to the annotated sequence on PomBase (https://www.pombase.org/, last accessed 03/06/2017). These differences resulted from the incorrect annotation of the position of the third intron (Figs. 1b, S1, S2). Additionally, we detected two splice variants (sv) of *hop1* (https://www.ebi.ac.uk/ena/data/view/LT963776; https://www.ebi.ac.uk/ena/data/view/LT907816). The main cDNA version (*hop1-sv1*) translated into a protein with 527 amino acids (AAs) in length which is one AA shorter than the currently annotated one (528 AAs). The discrepancy affects AAs 138-156/157 in the core region of the HORMA domain (Fig. 1). A second, shorter splice variant (*hop1-sv2*) seems to use a downstream AG as an alternative splice acceptor site at the third intron, thereby expanding it by another 21 bps (Figs. 1b, S1, S2). In a total of 25 cDNA clones from meiotic cDNA preparations of two independently constructed strains drawn from different time-points of a meiotic time-course, 23 were of the *hop1-sv1* type, one conformed to the *hop1- sv2* type, and one was aberrant representing a rather short form of hop1 with a frame-shift (Fig. 1c). Considering the scarcity of *hop1-sv2* in our samples, it could well represent a splicing accident or even a PCR artefact.

Using ClustalΩ (Sievers et al. 2014) the translation of the experimentally determined Hop1-sv1 HORMA domain sequence also produces a slightly better alignments with Hop1 HORMA domains of other *Schizosaccharomyces* (Fig. S3) species than the predicted one. The region of discrepancy in the experimentally derived *Sz. pombe* Hop1 HORMA domain (VCKFFNRLHESFIKLINVQKE) aligns without much gapping (only one AA position in *Sz. octosporus* and *Sz. cryophilus* shows a gap) and 7 fully conserved or strongly similar positions to its counterpart in the other *Schizosaccharomyces* species. The same region from the predicted *Sz. pombe* Hop1 (VSYQRSERFVIKLFLSGNVKTE) produces a large gap in the alignment with the *Sz. japonicus* sequence, receives a three AAs gap compared to *Sz. octosporus* and *Sz. cryophilus*, and aligns with only 4 fully conserved or strongly similar positions. Over the whole length of the HORMA domain in this multi-species comparison, this results in 10 fully conserved and 34 strongly similar AA residues in the alignment for the predicted *Sz. pombe* Hop1 sequence, and in 11 fully conserved and 35 strongly similar AA residues with the experimentally derived one (Fig. S3). This demonstrates that the current annotation of the *hop1* open reading frame on PomBase (https://www.pombase.org/) is erroneous, and that the long, predominant splice variant (*hop1-sv1*; https://www.ebi.ac.uk/ena/data/view/LT963776) we identified in our cDNA clones is the correct transcript of *hop1*.

### The long *hop1* splice variant *hop1-sv1* is sufficient for meiotic recombination

We were interested in establishing whether the long, predominant version of Hop1 (Hop1-sv1) is fully functional, and whether solely expressing the rare short version Hop1-sv2 (lacking 7 AAs in the HORMA domain, see Fig. 1) causes a phenotype. Additionally, we constructed a Hop1 version which lacks the HORMA domain altogether, Hop1-ΔHORMAD. In *S. cerevisiae* removing the N-terminal 268 AAs of Hop1 „ which includes the HORMA domain „ has previously been shown to abolish spore viability completely, however the C-terminal fragment remaining is still capable of binding DNA *in vitro* (Khan et al. 2013). In contrast, the *M127K* point mutation in the center of the HORMA domain of *C. elegans HTP-1*, one of four meiotic HORMA-paralogs in the nematode, results in only a minor meiotic phenotype despite HTP-1^M127K^ not localizing to meiotic chromosome axes (Silva et al. 2014). The *Sz. pombe* short form *hop1-sv2* is apparently not strongly expressed (if not a splicing accident or PCR artefact) in a wild-type background and thus unlikely to play a (major) role. However, in light of the distinct phenotypes of HORMA-domain mutants in budding yeast and *C. elegans*, we wanted to use the *hop1-sv2* variant and the *hop1-ΔHORMAD* mutant to probe the role of the HORMA domain for meiotic recombination and Hop1 localization to meiotic chromatin in fission yeast. The 7 AAs (positions 157-163) missing in Hop1-sv2 are close to the boundary between the HORMA core domain and the HORMA “safety-belt” region (Fig. 1a, b; Fig. S2) (Rosenberg and Corbett 2015; West et al. 2017), and their loss might affect interaction of Hop1 with the meiotic chromosome axis similar to nematode HTP-1^M127K^ and also interfere with Hop1’s self-oligomerization. The *hop1-ΔHORMAD* strains serve as controls, to monitor the effect removal of the complete HORMA domain has on meiotic recombination and Hop1 localization.

Previously, it has been shown that the full deletion of *hop1* causes a reduction in both intragenic (gene conversion) and intergenic (crossover) recombination at several genetic intervals in fission yeast with a concomitant increase in intrachromosomal recombination (Latypov et al. 2010). The *hop1* deletion also leads to defects in homologous pairing along chromosome I during meiosis (Latypov et al. 2010). These phenotypes can at least be partially explained by a reduction in DSB formation and a preponderance of DSBs being repaired from the sister chromatid rather than the homolog (Rothenberg et al. 2009; Latypov et al. 2010). Using our meiotic recombination assay system (Lorenz et al. 2010) which allows us to concomitantly measure spore viability, and the frequency of gene conversion (GC), crossovers (COs) and COs with GC events we found a statistically significant decrease in all recombination measures in a *hop1Δ* compared to a wild-type cross (Fig. 2). Indeed, GC decreased 4.7-fold (*p* = 4.53 × 10^−13^) (Fig. 2b), COs 2.8-fold (*p* = 3.61 × 10^−6^) (Fig. 2c), and COs associated with GC events were reduced by 8.7 percentage points (*p* = 0.01) (Fig. 2d) (see also Table S2). Overall, this reduction in recombination outcome did not negatively affect spore viability (Table S2). Recombination of a strain expressing only the long, predominant version *hop1-sv1* (from the *hop1*-promoter at its original locus without any marker genes inserted) was indistinguishable from wild type, whereas replacing the wild-type copy of *hop1* with *hop1-sv2* led to a moderate reduction in GC (1.4-fold, *p* = 1.51 × 10^−3^) and COs (1.7-fold, *p* = 2.27 × 10^−3^), but not in COs associated with GC events (*p* = 0.76) compared to wild type (Fig. 2, Table S2). The *hop1-ΔHORMAD* in turn was indistinguishable from a full deletion (Fig. 2, Table S2). These results indicate that in line with *hop1-sv1* being the predominant, if not the exclusive, splice variant behaves like wild-type *hop1* containing all its introns, that *hop1-sv2* lacking 21 nucleotides in the HORMA domain is not fully functional producing a hypomorphic phenotype, and that removal of the complete HORMA domain (*hop1-ΔHORMAD*) produces a null phenotype.

**Figure 2.**
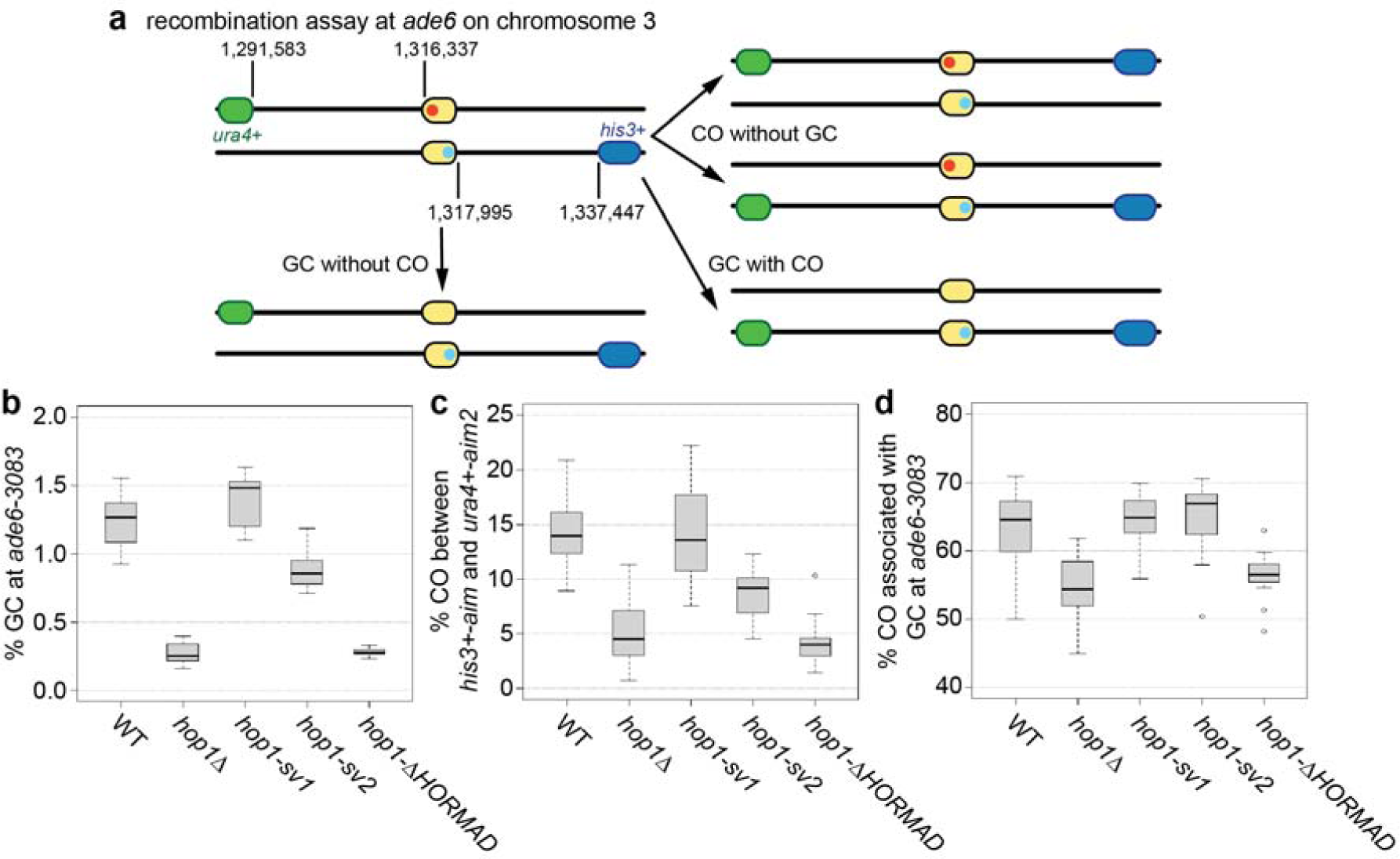
Wild-type and *hop1* mutant levels of meiotic recombination at the *ade6-3083* hotspot. (**a**) Schematic of the recombination assay at *ade6* (yellow) and its prevalent outcomes: positions of *ade6* and the artificially introduced markers *ura4^+^-aim2* (green) and *his3^+^-aim* (blue) on chromosome III are given in bp; positions of the point mutations in ade6 in the two parental strains are indicated in red (*ade6-3083*) and light blue (*ade6-469*). (**b**) Frequency of gene conversion (GC) at *ade6-3083*, (**c**) frequency of crossovers (COs) between *ura4^+^-aim2* and *his3^+^-aim*, and (**d**) percentage of COs between *ura4^+^-aim2* and *his3^+^-aim* associated with a GC event at *ade6-3083* in crosses of wild type (WT) ALP733×ALP731 (n = 12), *hop1Δ* UoA200×UoA199 (n = 17), *hop1-sv1* UoA766×UoA767 (n = 10), *hop1-sv2* UoA853×UoA854 (n = 12), and *hop1-ΔHORMAD* UoA945×UoA944 (n = 12); n indicates the number of independent crosses (see also Table S2).

### Hop1 localizes to meiotic chromatin and forms linear elements independent of a full-length HORMA domain

As with organisms possessing a fully-fledged synaptonemal complex, Hop1 localizes to meiotic chromosome axes, called linear elements, in fission yeast (Lorenz et al. 2004). However, the deletion of *hop1* has little impact on linear element formation per se (Lorenz et al. 2006). We were keen to understand whether the presence of the HORMA domain is required for localization of Hop1 to linear elements. We used immunostaining on chromatin spreads from meiotic fission yeast cells to detect heterozygously 13myc-tagged Hop1-sv1, Hop-sv2, and Hop1-ΔHORMAD in homozygous *hop1-sv1, hop1-sv2*, and *hop1-ΔHORMAD* strains, respectively. Hop1-sv1-13myc formed linear elements, and all four morphological classes of linear elements (dots, threads, networks, and bundles) were observed (Fig. 3a-d) (Bähler et al. 1993; Lorenz et al. 2004, 2006). This result is consistent with the wild-type phenotype for meiotic recombination of *hop1-sv1* crosses. Intriguingly, Hop1-sv2-13myc was indistinguishable from Hop1-sv1-13myc (Fig. 3e-h). Therefore, the loss of the 7 AAs close to the boundary between the HORMA core domain and the HORMA “safety-belt” region in the *hop1-sv2* allele does not conspicuously influence recruitment of Hop1 to linear elements, despite affecting meiotic recombination (Fig. 2b-d). Hop1-ΔHORMAD-13myc could not be detected on meiotic chromatin (Fig. S4) (a GFP-tagged version of the linear element protein Mug20 served as control to identify meiotic nuclei containing linear elements; (Estreicher et al. 2012)), this is consistent with *hop1-ΔHORMAD* displaying a meiotic recombination phenotype similar to *hop1Δ* (Fig. 2b-d).

**Figure 3.**
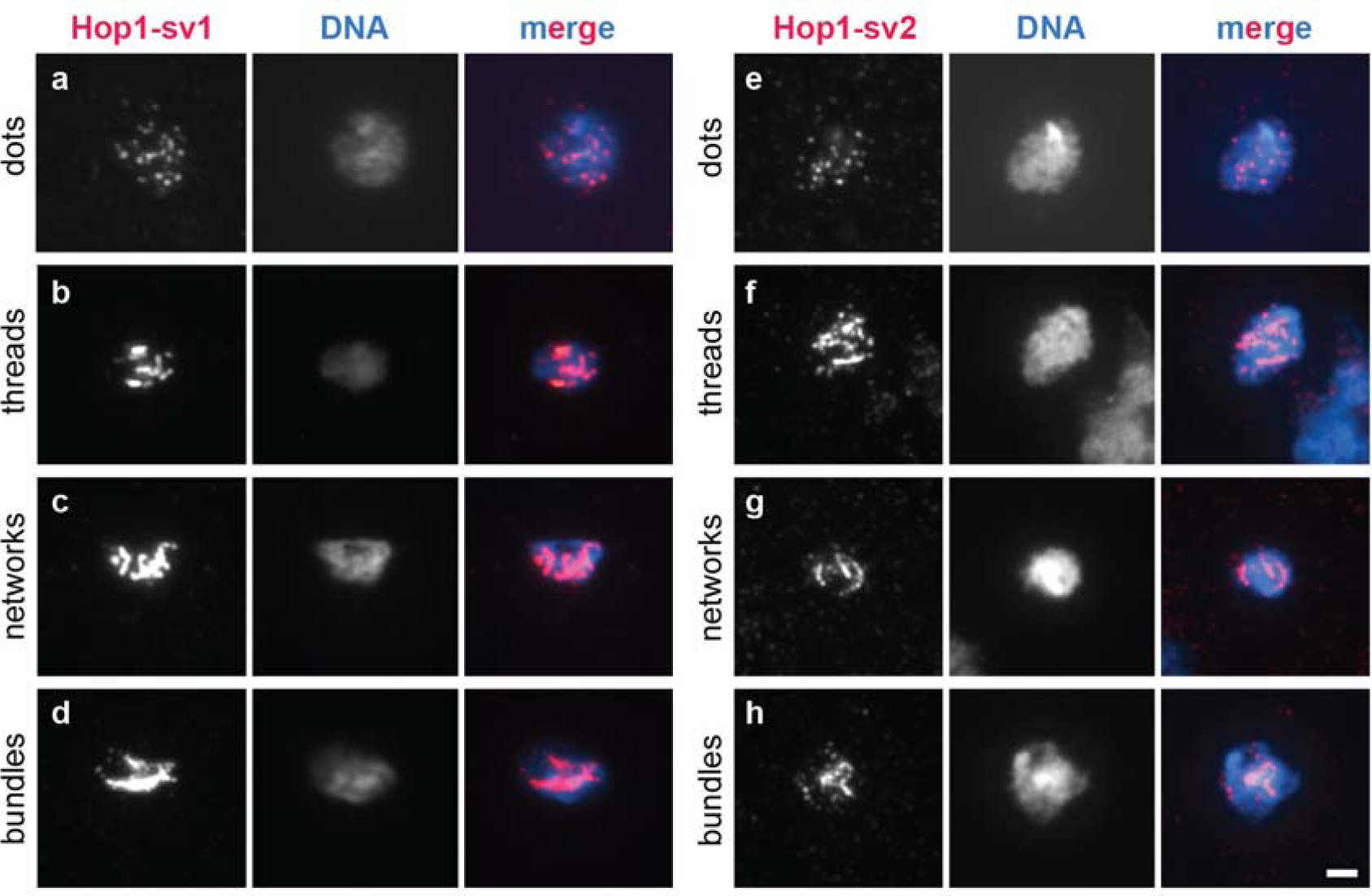
Linear element morphology delineated by immunostaining of Hop1-13myc on chromatin spreads from meiotic fission yeast cells. Linear elements as seen by immunostaining against Hop1-13myc develop from dots via threads to networks and bundles in *h*^*+N*^*/h*^-*smt0*^ *hop1-sv1/hop1-sv1::13myc-kanMX6 ade6-M210/ade6-M216* (UoA878) (**a-d**) or *h*^*+N*^*/h*^*-smt0*^ *hop1-sv2/hop1-sv2::13myc-kanMX6 ade6-M210/ade6-M216* (UoA930) (**e-h**). Hop1-13myc is shown in red and Hoechst 33342-stained DNA in blue in the merge panels. Scale bar represents 2 μm.

### Searching for Hop1 interactors

A bio-informatics screen using the *Pombe* Interactome (PInt) (Pancaldi et al. 2012) for potential Hop1 interactors retrieved *SPAC688.03c* as the highest hit from an uncharacterized open reading frame (10^th^ highest overall). The gene product of *SPAC688.03c* localizes preferentially to the nucleus (Matsuyama et al. 2006) (Fig. S5) and is upregulated at the onset of meiosis (Mata et al. 2002), which is consistent with a potential role in meiotic chromatin metabolism. Notably, a point mutation in the human ortholog of *SPAC688.03c, AMMECR1*, is associated with Alport syndrome (A), mental retardation (M), midface hypoplasia (M), and elliptocytosis (E) (Vitelli et al. 1999; Andreoletti et al. 2017). Therefore, we investigated the phenotype of a *SPAC688.03c* (from here on *aho1*, for *AMMECR1-homolog*) deletion in vegetative and generative cells.

First, *aho1Δ* was assayed for changes in genotoxin sensitivity and meiotic recombination outcome. The wild-type and *aho1Δ* strains exhibited similar levels of resistance to the topoisomerase I-poison camptothecin (CPT), the alkylating agent methyl methanesulfonate (MMS), the ribonucleotide reductase blocker hydroxyurea (HU), and ultraviolet (UV) radiation (Fig. 4a). The deletion of aho1 in a wild-type or *hop1Δ* background did not affect spore viability (Fig. 4b). The meiotic recombination outcome of an *aho1Δ* single mutant was similar to wild type, and an *aho1Δ hop1Δ* double mutant was indistinguishable from a *hop1Δ* single mutant, except for a moderate 1.4-fold reduction in gene conversion (*p* = 0.034) (Fig. 4c-e, Table S2). These results indicate that under standard laboratory conditions *aho1^+^* is not required for DNA damage repair or meiotic recombination. Therefore, Aho1 is not an essential interactor of Hop1 because it does not mirror the phenotype of *hop1Δ* and we did not observe strong genetic interactions with *hop1Δ*.

**Figure 4.**
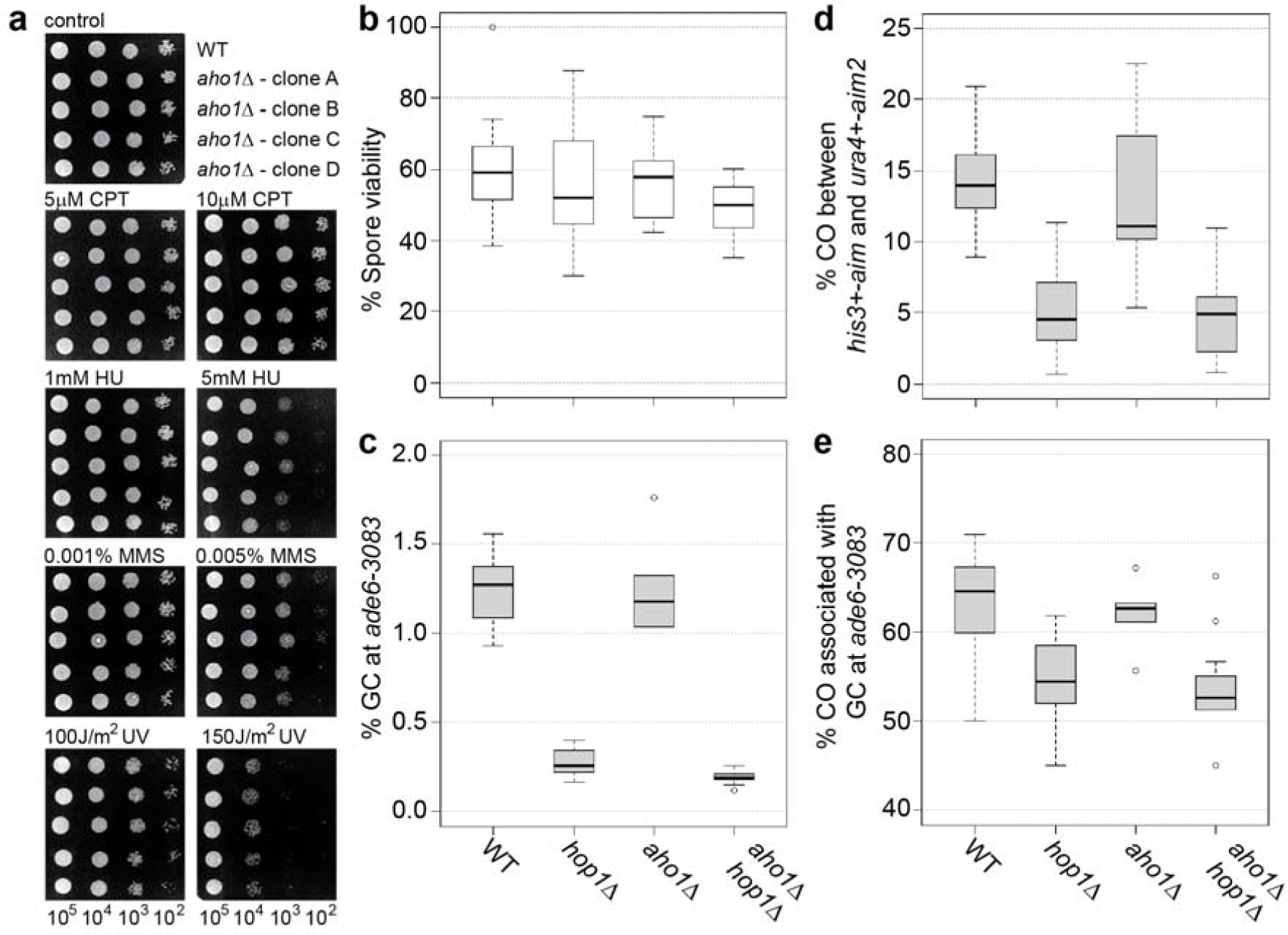
The role of *aho1^+^* in genotoxic stress resistance and meiotic recombination. (**a**) Spot assays comparing wild type (WT, FO652) and *aho1Δ* (UoA423) for sensitivity to a range of genotoxins; number of cells plated in each spot is indicated at the bottom of the figure. (**b**) Percent spore viability, (**c**) frequency of gene conversion (GC) at *ade6-3083*, (**d**) frequency of crossovers (COs) between *ura4^+^-aim2* and *his3^+^-aim*, and (**e**) percentage of COs between *ura4^+^-aim2* and *his3^+^-aim* associated with a GC event at *ade6-3083* in crosses of wild type (WT) ALP733×ALP731 (n = 12), *hop1Δ* UoA200×UoA199 (n = 16), *aho1Δ* UoA424×UoA427 (n = 6), and *aho1Δ hop1Δ* UoA785×UoA786 (n = 13); n indicates the number of independent crosses (see also Table S2).

We tested whether over-expression of *aho1^+^* might uncover a DNA repair and/or recombination role. The *aho1^+^* gene was put under the control of the thiamine-repressible *nmt1*-promoter at the medium pREP41-level and tagged with GFP at its C-terminus. Spot assays and recombination assays were performed after Aho1-GFP over-expression was induced for 24 hours (see also Fig. S5). Cells over-expressing Aho1-GFP did not show increased genotoxin sensitivities compared to cells containing an empty plasmid, cells over-expressing GFP alone, or cells grown in the presence of thiamine (in which expression is not induced) (Fig. 5a, b). Over-expression of Aho1-GFP also did not alter spore viability and meiotic recombination outcome in comparison to GFP over-expression (Fig. 5c-f, Table S3), indicating that a massive surplus of Aho1 protein has no negative impact on reproductive success.

**Figure 5.**
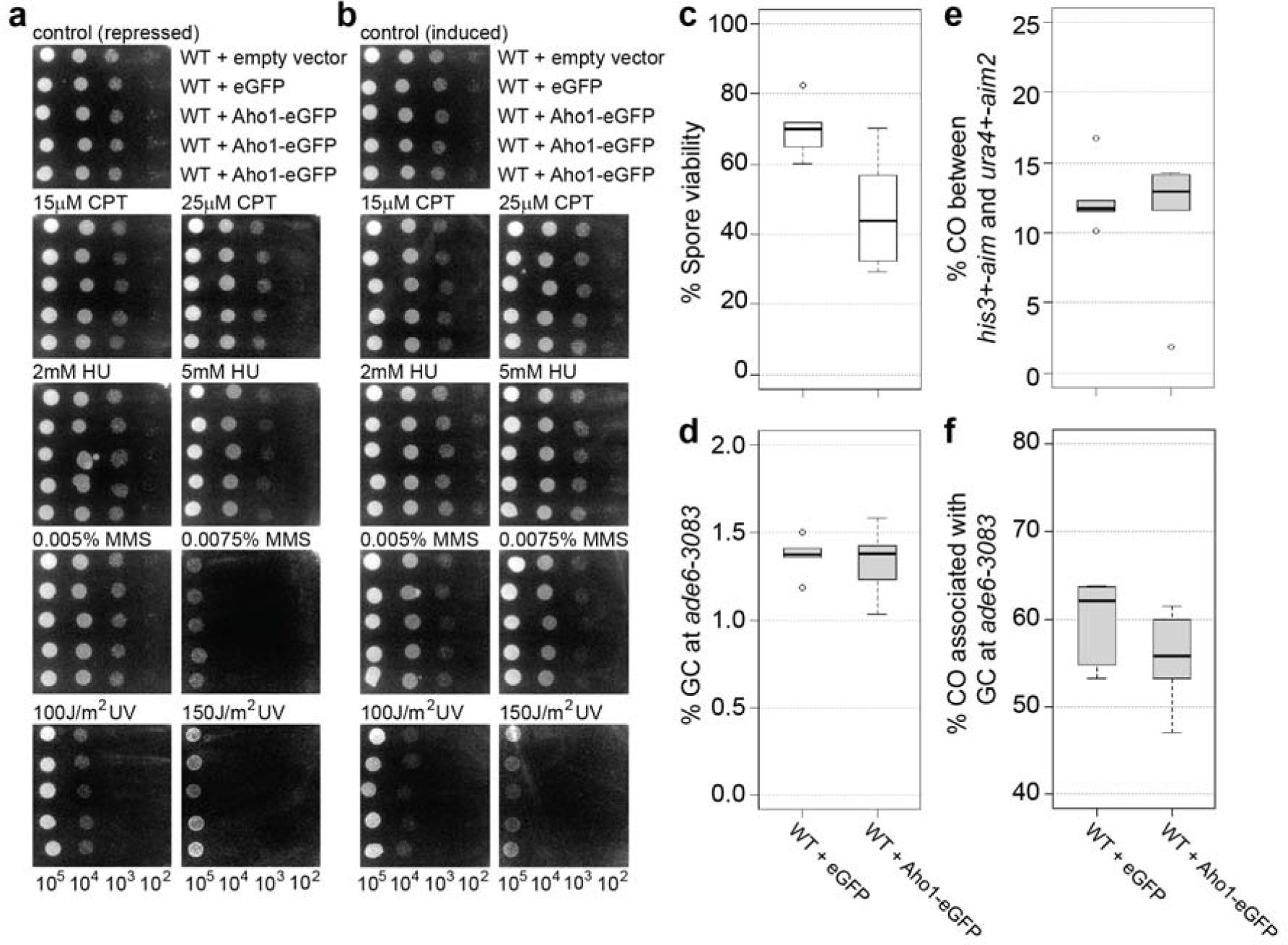
Over-expression of the *aho1^+^* in wild-type fission yeast cells and its consequences for genotoxic stress resistance and meiotic recombination. (**a, b**) Effect of GFP and Aho1-GFP over-expression on the genotoxin sensitivity of wild type (WT, FO652); number of cells plated in each spot is indicated at the bottom of the figure. (**c**) Percent spore viability, (**d**) frequency of gene conversion (GC) at *ade6-3083*, (**e**) frequency of crossovers (COs) between *ura4^+^-aim2* and *his3^+^-aim*, and (**f**) percentage of COs between *ura4^+^-aim2* and *his3^+^-aim* associated with a GC event at *ade6-3083* in crosses of wild type (WT) ALP733×UoA841 over-expressing GFP (n = 6), or Aho1-GFP (n = 6); n indicates the number of independent crosses (see also Table S3).

### *hop1* displays genetic interactions with the structure-selective endonuclease *mus81* and the DNA helicase *fml1*

Considering that the absence of *hop1* reduces recombination outcome in a similar fashion to Rad51/Dmc1-mediator (*rad55-rad57, swi5-sfr1*, and *rlp1-rdl1-sws1*) mutants (Lorenz et al. 2012, 2014) (Fig. 2) we were wondering how *hop1* genetically interacts with the key meiotic determinants of CO and non-CO formation, *mus81* and *fml1*.

The deletion of *hop1Δ* in a *mus81Δ* background improves spore viability from 1.85% to 18.7% (Fig. 6, Table S2) similar to what has been seen in *mus81Δ* Rad51/Dmc1-mediator double mutants (Lorenz et al. 2010, 2012, 2014). This rescue in spore viability is not accompanied by a restoration of CO frequency. A *hop1Δ mus81Δ* mutant shows only 0.2% of COs which is significantly different both from wild type (*p* = 1.9 × 10^−7^) and the *hop1Δ* single mutant (*p* = 2.7 × 10^−3^) (Fig. 6c, Table S2). Also, the rate of COs associated with GC events at 0.71% is significantly lower in the *hop1Δ mus81Δ* double mutant than in the wild type (63.0%, *p* = 5.3 × 10^−12^) or the *hop1Δ* single mutant (54.3%, *p* = 3.4 × 10^−15^) (Fig. 6d, Table S2).

**Figure 6.**
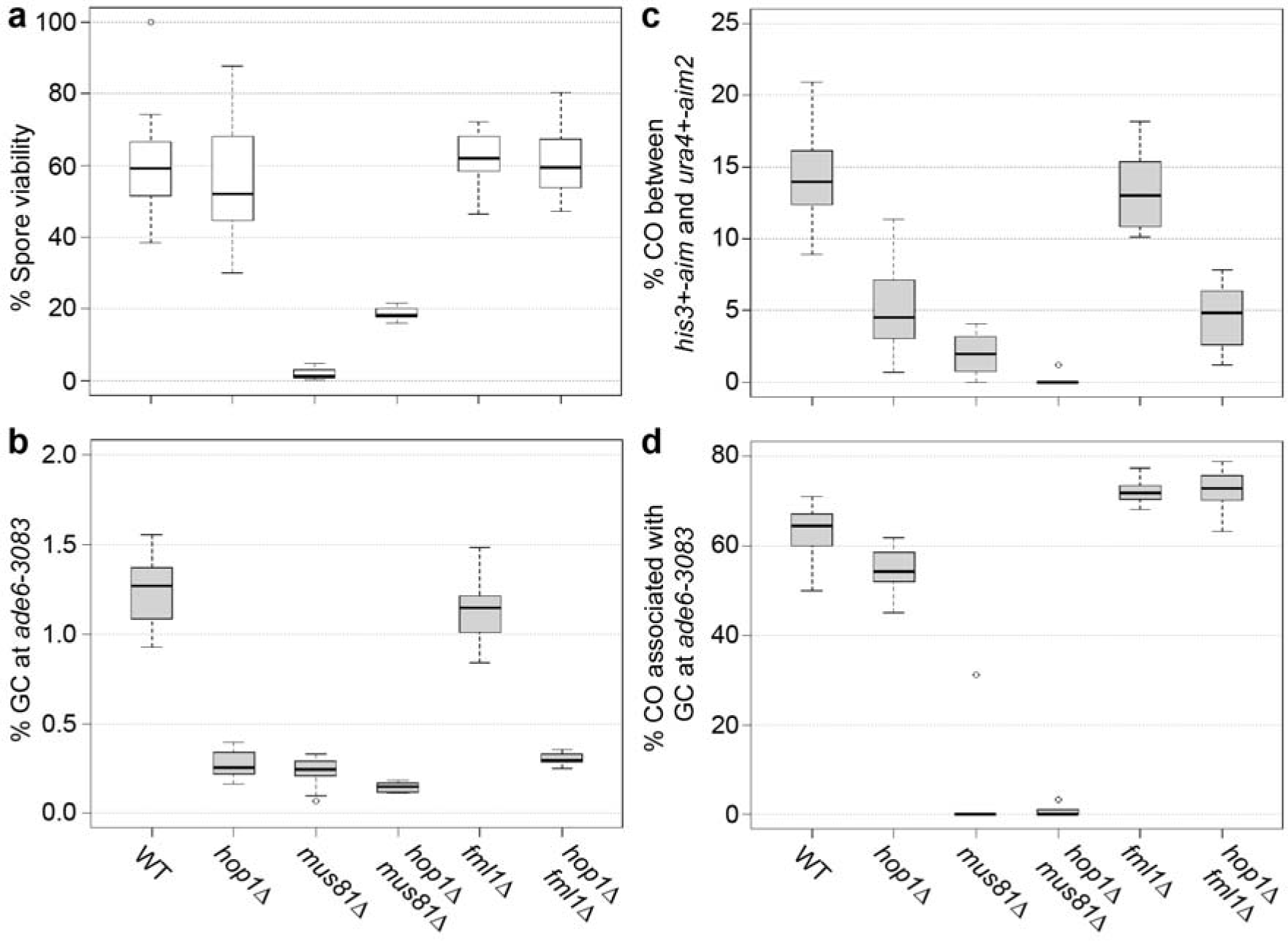
Genetic interaction of *hop1Δ* with deletions of factors required for meiotic recombination intermediate processing. (**a**) Percent spore viability, (**b**) frequency of gene conversion (GC) at *ade6-3083*, (**c**) frequency of crossovers (COs) between *ura4^+^-aim2* and *his3^+^-aim*, and (**d**) percentage of COs between *ura4^+^-aim2* and *his3^+^-aim* associated with a GC event at *ade6-3083* in crosses of wild type (WT) ALP733×ALP731 (n = 12), *hop1Δ* UoA200×UoA199 (n = 16), *mus81Δ* ALP812×ALP813 (n = 10; in a) and ALP802×ALP822 (n = 10; in c-d), *hop1Δ mus81Δ* UoA204×UoA203 (n = 6), *fml1Δ* ALP1133×FO2608 (n = 15), and *fml1Δ hop1Δ* UoA182×UoA181 (n = 14); n indicates the number of independent crosses (see also Table S2).

Removal of *fml1^+^* increases COs associated with GC events to 72.2% (~10 percentage points more than wild type) (Lorenz et al. 2012), but has little effect on other recombination outcomes (Fig. 6). In *fml1Δ hop1Δ* double mutants, there are 72.6% of COs associated with GC events which is significantly different from both wild type (*p* = 3.4 × 10^−3^) and a *hop1Δ* single mutant (*p* = 4.9 × 10^−9^), but indistinguishable from a *fml1Δ* single mutant (*p* = 0.78) (Fig. 6d, Table S2). Overall CO formation is not changed compared to *hop1Δ* (p = 0.78), but significantly different from wild type (*p* = 7 × 10^−7^) and *fml1Δ* (*p* = 6.3 × 10^−8^) (Fig. 6c, Table S2). Also GC frequency is unchanged compared to *hop1Δ* (*p* = 0.49) (Fig. 6b, Table S2) suggesting that in the absence of *hop1^+^* the DNA helicase Fml1 does not gain any additional functionality.

### *hop1* does not show strong genetic interactions with Rad51/Dmc1-mediator mutants

Rad51/Dmc1-mediators are highly conserved, positive determinants of meiotic recombination (Schwacha and Kleckner 1997; Ellermeier et al. 2004; Hayase et al. 2004; Bleuyard et al. 2005; Hyppa and Smith 2010; Lorenz et al. 2012, 2014; Sasanuma et al. 2013). In fission yeast the meiotic recombination phenotype of the *hop1Δ* crosses strongly resembles that of Rad51/Dmc1-mediator mutants in that it reduces all meiotic recombination outcomes and alleviates the effects of a *mus81Δ* mutants (see above). Therefore, we were interested in how *hop1* genetically interacts with the mediators *sfr1, rad55*, and *rlp1*, as well as *dmc1* itself. The deletion of *hop1^+^* leads to a notable reduction in DSB formation (Rothenberg et al. 2009; Latypov et al. 2010), and some of the decrease in recombination frequency in *hop1Δ* is likely due to low DSB formation. This reduction may be unrelated to Rad51/Dmc1-mediator function, because absence of the Sfr1-cofactor *swi5^+^* or of *dmc1^+^* does not appreciably reduce DSB formation (Young et al. 2004). We pose that in the absence of *hop1^+^*, which contributes to the establishment of a meiosis-specific chromatin environment, successful sexual reproduction will become more reliant on the mitotic repair factors Rad55 and Rlp1. Therefore, we expected a substantial negative genetic interaction of *hop1* with *sfr1, rad55, rlp1*, and *dmc1*.

Indeed, *hop1Δ rad55Δ* and *hop1Δ rlp1Δ* double mutants displayed significantly lower GC (*p* = 5.3 × 10^−5^ and *p* = 0.02, respectively) than a *hop1Δ* single mutant; the GC level is also notably lower than in *rad55Δ* (*p* = 2.9 × 10^−5^) and *rlp1Δ* (*p* = 1.3 × 10^−4^) (Fig. 7a, Table S2) (Lorenz et al. 2014). However, overall CO frequency and COs associated with GC events were not significantly different from *hop1Δ* or *rad55Δ* in *hop1Δ rad55Δ*, and from *hop1Δ* or *rlp1Δ* in *hop1Δ rlp1Δ* (Fig. 7b, c; Table S2). Deletion of *sfr1^+^* decreased GC rate by 7.5-fold in a *hop1Δ* background to 0.036% (*p* = 1.4 × 10^−8^) which is also significantly lower than the 0.13% (*p* = 1 × 10^−4^) found in an *sfr1Δ* single mutant (Fig. 7d, Table S2) (Lorenz et al. 2014). Other recombination outcomes were indistinguishable from *hop1Δ* (Fig. 7e, f; Table S2). The interaction of *hop1Δ* with *dmc1Δ* is completely epistatic (Fig. 7d-f, Table S2). Importantly, no noteworthy decreases of spore viability have been observed in any of the double mutants (Table S2). This stands in contrast to double mutant combinations of Rad51/Dmc1-mediators with each other or with *dmc1Δ*; *dmc1Δ rad55Δ, dmc1Δ rlp1Δ, rad55Δ sfr1Δ* and *rlp1Δ sfr1Δ* display strong synergistic reductions in spore viability (Lorenz et al. 2014). Overall, this suggests that in fission yeast meiosis the absence of *hop1^+^* does not generate a situation which necessitates the presence of a fully functional recombination machinery to produce viable progeny.

**Figure 7.**
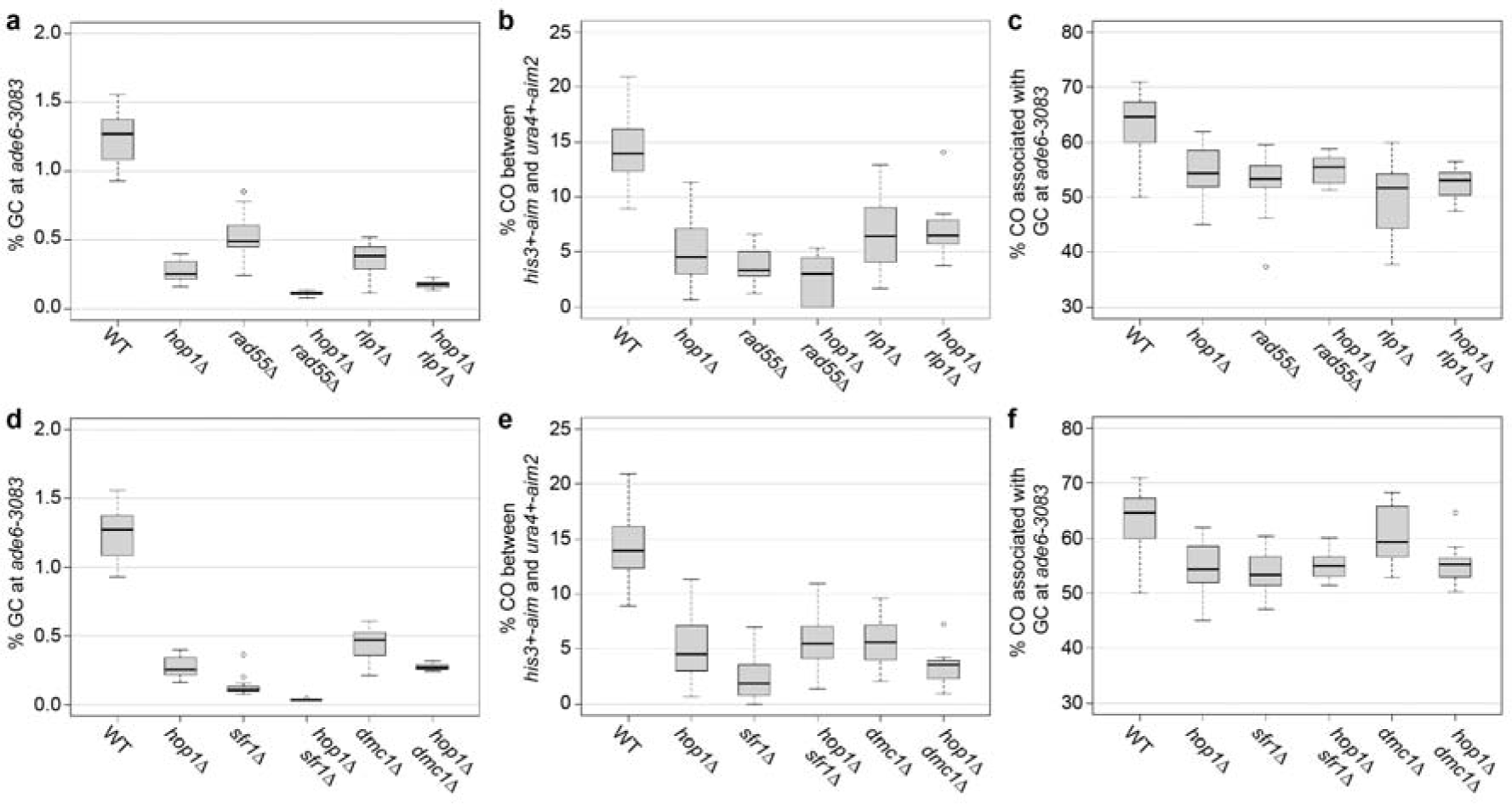
Genetic interaction of *hop1Δ* with deletions of Rad51/Dmc1-mediator genes. (**a,d**) frequency of gene conversion (GC) at *ade6-3083*, (**b, e**) frequency of crossovers (COs) between *ura4^+^-aim2* and *his3^+^-aim*, and (**c,f**) percentage of COs between *ura4^+^-aim2* and *his3^+^-aim* associated with a GC event at *ade6-3083* in crosses of wild type (WT) ALP733×ALP731 (n = 12), *hop1Δ* UoA200×UoA199 (n = 16), *rad55Δ* ALP1649×ALP1648 (n = 12), *hop1Δ rad55Δ* UoA846×UoA847 (n = 10), *rlp1Δ* ALP1623×ALP1620 (n = 18), *hop1Δ rlp1Δ* UoA848×UoA849 (n = 10), *sfr1Δ* ALP800×ALP782 (n = 22), *hop1Δ sfr1Δ* UoA844×UoA845 (n = 12), *dmc1Δ* ALP1545×ALP1544 (n = 12), and *dmc1Δ hop1Δ* UoA851×UoA852 (n = 10); n indicates the number of independent crosses (see also Table S2).

## Discussion

Here we characterize the genetic interactions of *hop1*, which encodes a meiotic HORMA domain protein, with homologous recombination factors in fission yeast. Meiotic HORMADs are components of the meiosis-specific chromosome axis which promote DSB formation to allow for the efficient pairing of homologous chromosomes. At the same time HORMADs attenuate undesirable recombination between sister chromatids (reviewed in Rosenberg and Corbett 2015). In organisms possessing a synaptonemal complex, HORMADs support its formation, and depending on the organism may mount a checkpoint arrest when DSB repair is not completed or homologous chromosomes are not synapsed (Woltering et al. 2000; Armstrong et al. 2002; Couteau and Zetka 2005; Martinez-Perez and Villeneuve 2005; Carballo et al. 2008; Daniel et al. 2011; Rinaldi et al. 2017). *Sz. pombe* meiosis progresses without the formation of a synaptonemal complex, therefore any roles related to the synaptonemal complex will likely not apply. Indeed, DSB formation is reduced and inter-sister recombination is increased in fission yeast *hop1Δ* (Rothenberg et al. 2009; Latypov et al. 2010). Consistent with DSB reduction, decreased numbers of Rec7-foci (an essential co-factor of Spo11) have been observed when *hop1* was absent (Lorenz et al. 2006). As a result of DSB reduction, meiotic inter-homolog recombination frequencies are lower in *hop1Δ* than in wild type (Latypov et al. 2010) (Fig. 2). Thus, the Hop1 function in promoting DSB formation seems to be conserved in fission yeast. In budding yeast Hop1 also acts downstream of DSB formation in promoting Dmc1-dependent inter-homolog recombination by preventing Dmc1-independent Rad51-driven recombination (Niu et al. 2005; Carballo et al. 2008). The suppression of inter-sister recombination is achieved by Hop1-dependent activation of Mek1 which in turn phosphorylates and deactivates the Rad51-mediator Rad54, and also phosphorylates and activates the meiosis-specific Rad51-inhibitor Hed1 (Tsubouchi and Roeder 2006; Niu et al. 2009; Callender et al. 2016). Together, this regulatory network has specific genetic consequences: (I) meiotic recombination cannot proceed without Dmc1 which causes a meiotic arrest resulting in an inability to sporulate, and (II) deleting *hop1* in a *dmc1Δ* background restores spore formation by allowing Rad51-dependent DSB repair (Niu et al. 2005; Carballo et al. 2008). In fission yeast the genetic interaction between *hop1* and *dmc1* looks quite different (Fig. 7) in that it is epistatic, however neither single mutant has as strong a phenotype as the corresponding mutants in *S. cerevisiae* (Hollingsworth and Byers 1989; Bishop et al. 1992; Grishchuk and Kohli 2003; Latypov et al. 2010). Two features may account for this apparent difference between the two yeasts: The budding yeast is probably the only organism which shuts down Rad51 strand exchange activity via Hed1 (Tsubouchi and Roeder 2006), and fission yeast employs both Rad51 and Dmc1 in their direct strand exchange capacity (Grishchuk and Kohli 2003). Interestingly, the genetic interaction of meiotic *Hormad* and *dmc1* genes generally does not show much conservation. In *Arabidopsis* mutation of *asy1* or *dmc1* causes a strong reduction or absence of chiasmata, respectively, leading to fertility defects. However, the *asy1 dmc1* double mutant looks like a dmc1 single mutant (Sanchez-Moran et al. 2007). In the mouse both *Dmc1^-/-^* and *Hormad1^-/-^* single mutants undergo meiotic arrest at mid-pachytene, and so does the *Dmc1^-/-^ Hormad1^-/-^* double mutant (Daniel et al. 2011), i.e. there is no reciprocal rescue as in *S. cerevisiae hop1Δ dmc1Δ*. These disparate observations in various model organisms suggest that the genetic networks controlling DSB repair and CO formation, and how these processes are integrated with chromosome axis organization, have undergone considerable rewiring during evolution. Because fission yeast relies on both Rad51 and Dmc1 to invade a repair template during meiotic DSB repair, we also looked at the genetic interaction of *hop1* with Rad51/Dmc1-mediator mutants. Apart from an additive effect on GC frequency in *hop1Δ rad55Δ, hop1Δ rlp1Δ*, and *hop1Δ sfr1Δ* double mutants, which can easily be explained by the early functions of Hop1 (reduced DSB formation and increased inter-sister recombination), other recombination outcomes were not different from the *hop1Δ* single mutant (Fig. 7). This indicates that in a fission yeast *hop1Δ* mutant gamete production does not become more reliant on strand exchange than in a wild-type background.

Genetic interaction of *hop1Δ* with *mus81Δ* resembled that of *mus81Δ* combined with mutants affecting strand exchange (Ellermeier et al. 2004; Lorenz et al. 2012, 2014), rescuing *mus81Δ*’*s* spore viability defect without rescuing its CO defect (Fig. 6). This interaction can be explained by Hop1 promoting DSB formation and inter-homolog recombination: Fewer DSBs and fewer inter-homolog events in *hop1Δ* decrease the probability of the production of recombination intermediates requiring the action of Mus81. Additionally, the absence of *hop1* might allow DNA helicases to process recombination intermediates as non-COs more efficiently than in a wild-type background.

It is well established that fungal Hop1 localizes to the chromosome axis during meiotic prophase by physically interacting with the linear/lateral element protein Rec10/Red1 in fission and budding yeast, respectively (de los Santos and Hollingsworth 1999; Spirek et al. 2010). Considering Hop1’s central role in coordinating meiotic progression, it is not unlikely that it would physically interact with previously uncharacterized meiotic players; indeed with IHO1, a novel HORMAD1-interactor has recently been described in mice (Stanzione et al. 2016). A bio-informatical protein interaction-predictor PInt (Pancaldi et al. 2012) suggested that the uncharacterized AMMECR1-homolog Aho1 might be a potential Hop1-interactor in fission yeast. However, neither deletion nor over-expression of Aho1 indicates a clear meiotic function, independent of the presence or absence of Hop1 (Figs. 4, 5).

One of the determining features of Hop1 is its HORMA domain whose function was enigmatic until recently, when it was implicated in the recruitment of Hop1 to the meiotic chromosome axis and in its self-assembly into larger homomeric complexes (Kim et al. 2014; Rosenberg and Corbett 2015). We exploited a cDNA clone of *hop1* (*hop1-sv2*) which produces a protein that lacks 7 AAs in the HORMA domain and constructed a version of hop1 lacking the complete HORMA domain (*hop1-ΔHORMAD*) to examine whether the HORMA domain in fission yeast Hop1 is essential for its function and localization. Interestingly, expressing Hop1-sv2 in place of wild-type Hop1 did not result in a notable reduction in localization to chromatin, however it did cause a hypomorphic phenotype for GC and CO frequency outcome, whereas the complete removal of the HORMA domain from Hop1 mimicked a deletion phenotype (Figs. 2, 3, S4). This suggests that the shortened HORMA domain in Hop1-sv2 is still partially functional, because absence of Hop1’s HORMA domain in fission yeast completely abolishes localization to meiotic chromatin causing a *hop1Δ* phenotype in recombination outcome.

It is very interesting how genetic pathways driving meiotic chromosome organization and meiotic recombination are rewired during evolution. The fission yeast, which is lacking a synaptonemal complex, but retains remnants of a meiosis-specific chromosome axis, is a great model for probing the roles of conserved meiotic factors such as Hop1. Clearly, Hop1 fulfills an important meiotic function in promoting DSB formation and inter-homolog recombination in this organism, but lacks regulatory roles in lieu of the synaptonemal complex, which appears crucial in the majority of organisms performing meiosis with a synaptonemal complex.

## Acknowledgments

We are grateful to Shin-ichiro Hiraga, Franz Klein, Jürg Kohli, Josef Loidl, Walter W. Steiner, Matthew C. Whitby, and the National BioResource Project (NBRP) Japan for providing strains and/or plasmids, and to K.C. Chan, M. Roca, and D. Whyte for technical assistance. Kevin Corbett, Nancy Hollingsworth, Takashi Kubota, and Josef Loidl are thanked for critically reading a previous version of this manuscript. Microscopy was performed at the University of Aberdeen Microscopy & Histology facility (Kevin Mackenzie). This work was supported by the Biotechnology and Biological Sciences Research Council UK (BBSRC, Doctoral Training Grant BB/F016964/1), and the University of Aberdeen (College of Life Sciences and Medicine Start-up grant to AL).

## Author contributions

SDB: unpublished essential reagents (yeast strains, plasmids), acquisition of data, analysis and interpretation of data, revising the manuscript; ODJ: acquisition of data, revising the manuscript; AL: conception and design, unpublished essential reagents (yeast strains, plasmids), acquisition of data, analysis and interpretation of data, drafting and revising the manuscript.

